# Short term supplementation of celecoxib shifted butyrate production on a simulated model of the gut microbial ecosystem and ameliorated *in vitro* inflammation

**DOI:** 10.1101/679050

**Authors:** Emma Hernandez-Sanabria, Evelien Heiremans, Marta Calatayud Arroyo, Ruben Props, Laurent Leclercq, Jan Snoeys, Tom Van de Wiele

## Abstract

Celecoxib has been demonstrated effective in the prevention and treatment of chronic inflammatory disorders through inhibition of altered cyclooxygenase-2 (COX-2) pathways. Despite the benefits for preventing colorectal cancer (CRC), continuous administration may increase risk of cardiovascular events. Understanding microbiome-drug-host interactions is fundamental for improving drug disposition and safety responses of colon-targeted formulations, but little information is available on the bidirectional interaction between individual microbiomes and celecoxib. Here we conducted *in vitro* batch incubations of faecal microbiota to evaluate the short-term impact of celecoxib on activity and composition of colon bacterial communities. Celecoxib-exposed microbiota shifted metabolic activity and community composition, whereas total transcriptionally active bacterial population was not significantly changed. Butyrate production decreased by 50% in a donor-dependent manner, suggesting that celecoxib impacts *in vitro* fermentation. Microbiota-derived acetate has been associated with inhibition of cancer markers and our results suggest uptake of acetate for bacterial functions when celecoxib was supplied, which potentially favoured bacterial competition for acetyl-CoA. We further assessed whether colon microbiota modulates anti-inflammatory efficacy of celecoxib using both a simplified inflammation model, and a novel *in vitro* simulation of the enterohepatic metabolism. Celecoxib was responsible for only 5% of the variance in bacterial community composition but celecoxib-exposed microbiota preserved barrier function and decreased concentrations of IL-8 and CXCL16 in a donor-dependent manner in our two cell models simulating inflammatory milieu in the gut. Our results suggest that celecoxib-microbiome-host interactions may not only elicit adaptations in community composition but also in microbiota functionality and may need to be considered for guaranteeing efficient COX-2 inhibition.

**IMPORTANCE:** As inter-individual changes in the microbiome composition and functionality may be a confounder on pharmacotherapy, we obtained mechanistic understanding on how short-term celecoxib exposure impacts the functional activities of colon communities. Celecoxib-exposed microbiota shifted metabolic activity without impacting numbers of total active bacteria, but only community composition. Thus, increased relative abundance of particular genera during celecoxib supplementation may just indicate changes in maintenance energy. Focus on the influence of acetyl-CoA on cancer cells and verifying whether changes in acetate:propionate:butyrate ratios rather than in taxonomic diversity can be used as markers of decreased inflammation may be the next frontiers for predicting successful NSAID therapy, and ultimately for developing microbiome-based therapies.

## INTRODUCTION

Colorectal cancer (CRC) is the second most common cancer in the developed world (1), and it is expected to increase by 60% in the next 20 years (2). Increased synthesis of prostaglandins is a feature of chronic inflammation and has been proposed to promote tumorigenesis in the colon (3). Prostaglandin E_2_ (PGE_2_) is a key mucosal inflammatory mediator produced when membrane phospholipids are metabolized by cyclooxygenase (COX) enzymes. Previous research showed that COX-2 expression was increased in CRC (4), causing proliferation, enhanced angiogenesis and apoptosis suppression (5). In this way, COX-2 inhibitors such as celecoxib are among the most promising chemopreventive agents for CRC (6). Despite these benefits, their long-term use may shift cardiovascular homeostasis, inhibiting biosynthesis of vascular COX-2 dependent prostacyclin (PGI_2_) (7) and increasing cardiovascular events risk (8).

Advances in omics techniques have provided insights on the role of the gut microbiome in cancer development (9). Currently, the driver-passenger model suggests that intestinal “driver” bacteria contribute to initiate CRC by damaging epithelial DNA (10). This microenvironment alteration impairs barrier function and favours opportunistic “passenger” bacteria, placing selective pressure on the microbiome (10).

When colon-targeted drugs are orally administered, they encounter the gut microbiota before reaching host tissues. Microbial drug metabolism may impact drug stability and activity, influencing toxicity and inflammation on the host (11). Conversely, drugs may also alter the gut microbiota, modulating community structure, and potentially resulting in dysbiosis (12). Understanding microbiome-drug-host interactions will be crucial to assess the potential impact on ADME and safety responses of intestinal targets. Although the gut microbiome plays an enormous role in patient health, and in pharmacokinetic and pharmacodynamic response (13), pharmaceutical companies have only recently started to explore the microbiome as potential contributor to drug outcomes. Similarly, regulatory agencies are not currently considering gut bacteria within the official evaluations before approval of colon-targeted drugs. Besides the microbial modulation of drug bioavailability and pharmacokinetics (11), low solubility and the small aqueous volume in the colon may additionally hinder drug efficacy (14). As interest in developing novel colon-targeted drug delivery systems for CRC has increased, insight into the presystemic biotransformation by the gut microbiota is fundamental. *In vitro* studies have shown that bacteria can metabolize celecoxib (15), potentially jeopardizing its anti-inflammatory effects. Additionally, whether celecoxib presence can modulate the host-microbiome crosstalk remains unresolved. Prior reports correlating gut microbiome changes with celecoxib treatment relied on DNA amplicon sequencing (16). As the presence of dead bacterial cells may obscure the interpretation from microbiome data, we analysed the bacterial community composition based on cDNA, as a proxy for the metabolically active community.

Colonic transit times (17-24 hrs) (17, 18) are longer compared to those of the small intestine (3-4 hrs), and overall systemic absorption of poorly soluble drugs may be improved with colon-delivery systems (17). Although celecoxib can be absorbed throughout the GI tract, we simulated delivery in a proximal colon-targeted formulation because of the environment vicinity between proximal colon and small intestine. Thus, the aim of this research was to screen the short-term bidirectional interactions between celecoxib and the microbiome of the *in vitro* proximal colon, and to elucidate the ultimate impact of the microbiota on the anti-inflammatory efficacy of celecoxib.

## RESULTS

### Celecoxib impacts bacterial fermentative metabolism in a donor-dependent manner

Short-chain Fatty Acids (SCFAs) were used to monitor functional activities of the colon communities following exposure to celecoxib (CX) or its vehicle (PEG). Total SCFA production from separate faecal microbiota incubations of 8 different individuals was significantly lower for CX incubations compared to those with PEG (*P* < 0.05, Table 1), but pH remained stable throughout the experiment (Table 1). While the 60:20:20 ratio of the three major SCFA (acetate, propionate, and butyrate) was consistent across donors at the initial time point (Supplementary Table 1A), a shift was evident at the end of the incubation. Both relative and absolute concentrations of butyrate and propionate were higher in samples treated with only PEG compared to those with CX. In fact, celecoxib significantly lowered butyrate after 16 hours (*P* < 0.05, Table 1), except in donors 1, 4 and 8 (Supplementary Table 1B). In contrast, total active bacterial load remained unchanged when celecoxib was supplemented (Table 1, Supplementary Table 2). These findings might indicate that bacteria either alter their metabolic activity when exposed to CX or that changes in bacterial composition lead to shifts in fermentation profiles.

**Table 1.** Bacterial functionality metrics following short-term incubation with either celecoxib or carrier (n = 8). CX, celecoxib; PEG, carrier. SEM, standard error of the mean.

| Variable | Treatment<br>(Mean ± SEM) | | *P* value |
| --- | --- | --- | --- |
|  | CX | PEG |  |
| Acetate (mM) | 11.51 ± 1.44 | 13.31 ± 1.50 | 0.39 |
| Propionate (mM) | 3.29 ± 0.41 | 3.86 ± 0.42 | 0.33 |
| Butyrate (mM) | 1.57 ± 0.33 | 3.20 ± 0.33 | 0.0007 |
| Valerate (mM) | 0.61 ± 0.15 | 0.78 ± 0.15 | 0.45 |
| Isobutyrate (mM) | 0.30 ± 0.07 | 0.34 ± 0.07 | 0.70 |
| Isovalerate (mM) | 2.70 ± 0.52 | 3.30 ± 0.54 | 0.42 |
| Caproate (mM) | 0.15 ± 0.04 | 0.12 ± 0.04 | 0.61 |
| Isocaproate (mM) | 0.001 ± 0.002 | 0.01 ± 0.002 | 0.009 |
| Total SCFA (mM) | 20.14 ± 2.37 | 24.91 ± 2.45 | 0.16 |
| Total Branched SCFA (mM) | 2.56 ± 0.49 | 3.25 ± 0.52 | 0.34 |
| St:Br ratio | 17.76 ± 4.48 | 12.71 ± 4.31 | 0.42 |
| % of Acetate | 60.87 ± 2.60 | 53.58 ± 2.77 | 0.06 |
| % of Propionate | 16.17 ± 0.65 | 15.75 ± 0.70 | 0.67 |
| % of Butyrate | 6.57 ± 0.64 | 12.99 ± 0.68 | <0.0001 |
| Acetate:Propionate:Butyrate ratio | 61:16:7 | 54:16:13 | - |
| pH | 5.76 ± 0.04 | 5.75 ± 0.04 | 0.86 |
| Total copy number of 16S rRNA gene | 3.37E09 ± 3.33E08 | 4.35E09 ± 3.33E08 | 0.04 |

### Bacterial abundance remains stable, but microbiome diversity and composition shifted following celecoxib supplementation

Total DNA may also include that of dead bacterial cells, so we analysed community composition based on cDNA, as a proxy for the metabolically active bacterial community Quantitative RT-PCR revealed that total 16S rRNA gene copy numbers from metabolically active bacteria were not significantly different between treatments, across donors (Supplementary Table 2). In contrast, community composition and structure were influenced, as celecoxib was associated with reduced diversity at the end of the incubation (Figure 1, Table 2A and 2B), with significant donor effect (PERMANOVA *P* < 0.001, Figure 2). CX shifted alpha diversity and community composition metrics across donors (Supplementary Table 3, Figure 3). Although interindividual variability was significant, metabolically active taxa with the highest relative abundances in control incubations were: *Bacteroides* sp., *Faecalibacterium* sp., Ruminococcaceae UGC002, and *Megasphaera* sp. In contrast, *Alistipes* sp., *Dialister* sp., *Escherichia*-*Shigella* sp., *Lachnospira* sp., *Megamonas* sp., *Parabacteroides* sp., *Roseburia* sp., and *Sutterella* sp. were highly abundant when CX was supplied (Figure 3).

**Figure 1.**
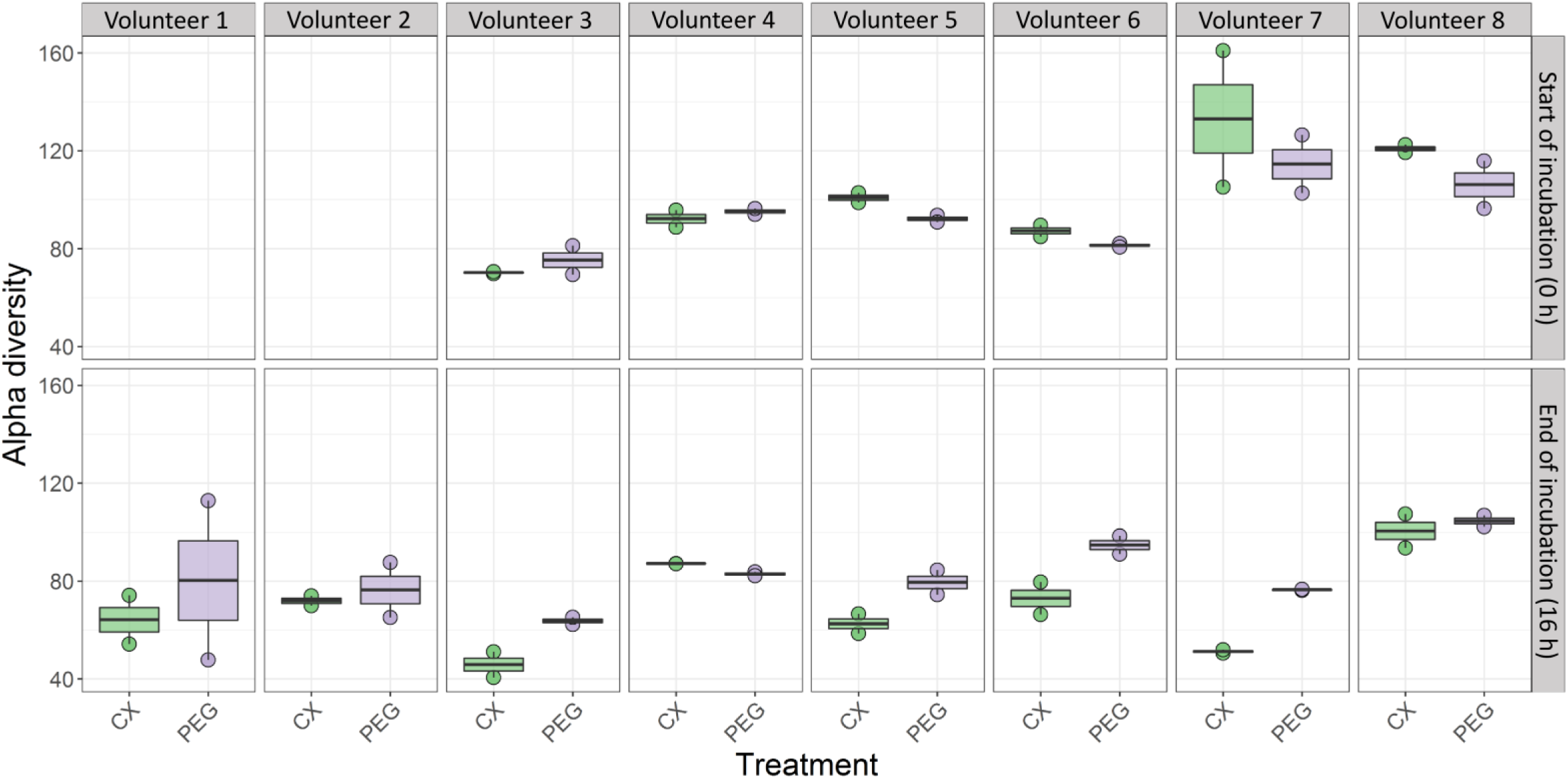
Inverse Simpson index (*D*_2_) was used to assess alpha diversity dynamics at the beginning and at the end of the incubation (0 and 16 h post treatment) on each volunteer. Horizontal bars indicate means and distribution of each data point is indicated. Alpha diversity for celecoxib treatment (CX) is indicated in green boxes, while purple is for this metric observed when the control treatment with the vehicle (PEG) was supplied.

**Figure 2.**
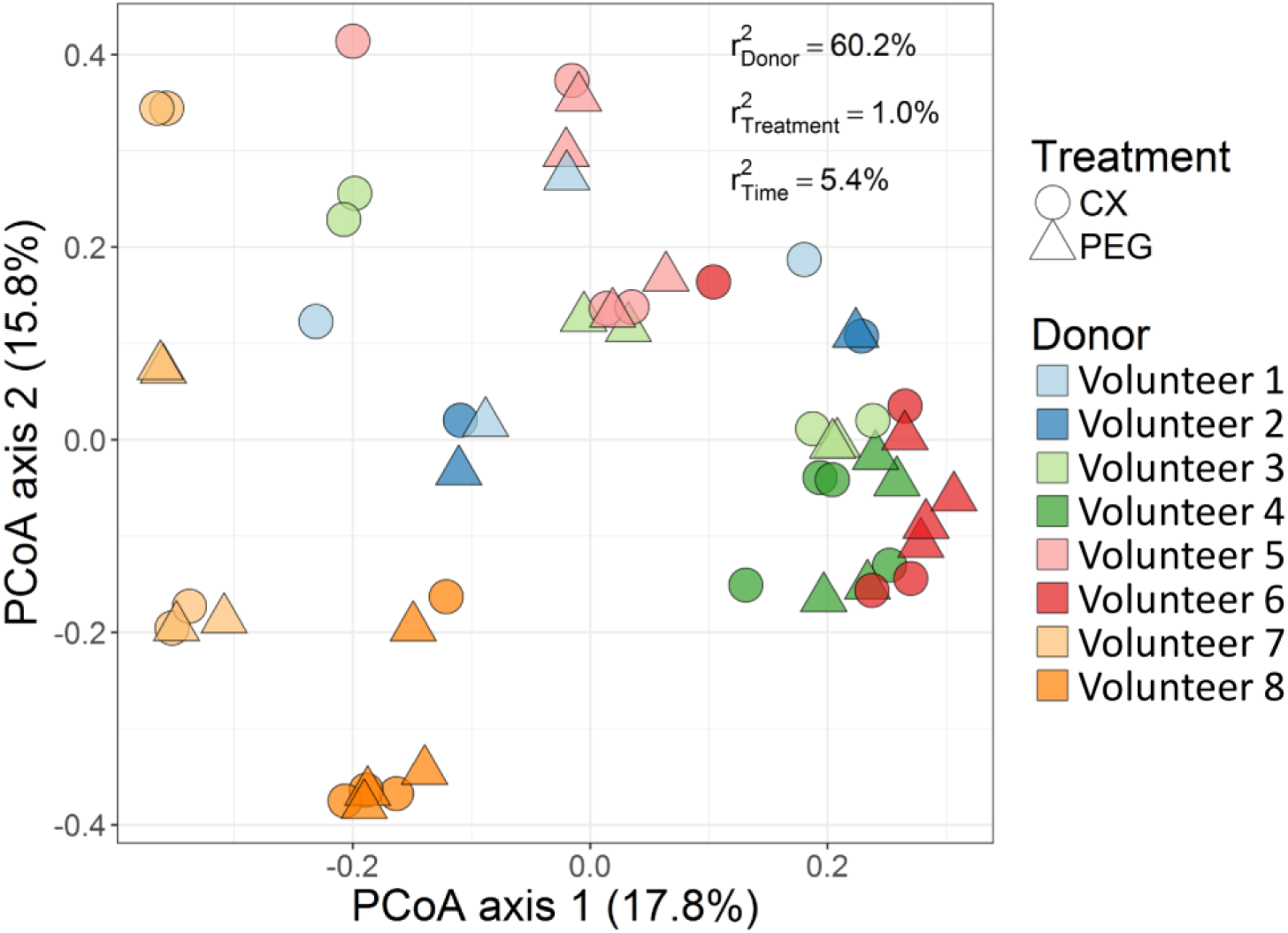
Interindividual differences drive bacterial community composition. Principal coordinate analysis (PCoA) revealed differences in community structure between volunteers. The variance explained by the experimental factors (*P* < 0.01) was calculated using PERMANOVA analysis and it is indicated on the top right. Donor was the factor significantly contributing to the variations among individual microbiomes

**Figure 3.**
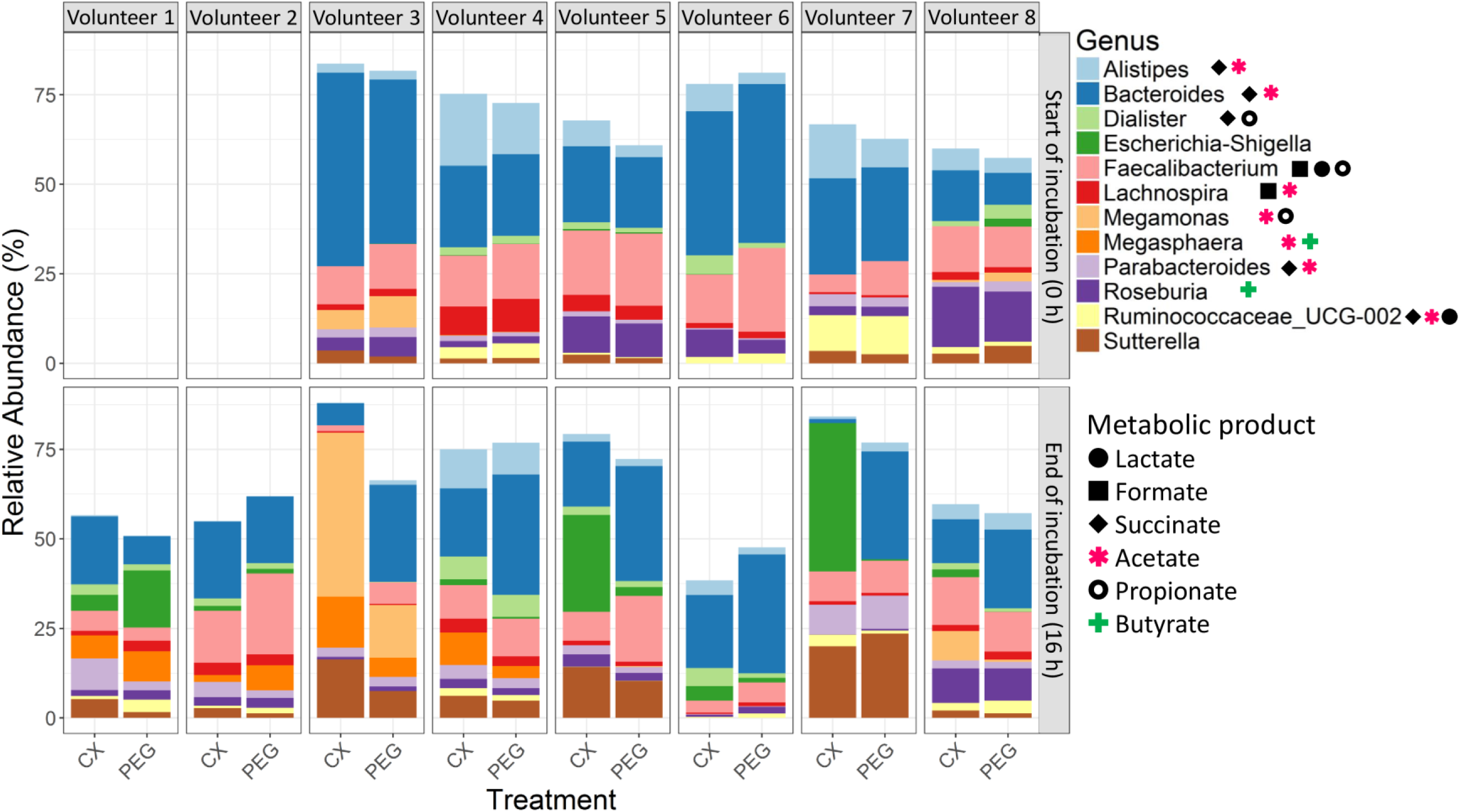
Relative abundances of the bacterial genera occurring when celecoxib (CX) or a control treatment (PEG) were supplied. The upper panel describes the community composition at the beginning of the treatment (0 h), while the lower panel presents the 12 taxa with the highest activity-based (cDNA-based) relative abundances after 16 h of treatment.

**Table 2A.** Impact of donor and supplementation of celecoxib (CX) or vehicle (PEG) on initial bacterial community metrics (time 0 h), n = 8.

| Metric | Treatment |  | SEM | P value |  |  |
| --- | --- | --- | --- | --- | --- | --- |
|  | CX | PEG |  | Treatment | Donor | Treatment*Donor interaction |
| Evenness | 0.61 | 0.60 | 0.0009 | 0.83 | 0.0001 | 0.23 |
| Shannon ($H'$) | 2.71 | 2.72 | 0.03 | 0.81 | <0.0001 | 0.13 |
| Simpson | 0.85 | 0.85 | 0.007 | 0.43 | <0.0001 | 0.04 |
| Inverse Simpson ($D_2$) | 8.00 | 8.48 | 0.53 | 0.54 | <0.0001 | 0.76 |
| Fisher's diversity | 13.41 | 13.16 | 0.25 | 0.49 | <0.0001 | 0.34 |
| Total Richness (S) | 92.50 | 95.83 | 3.83 | 0.55 | 0.002 | 0.77 |
| Chao | 144.12 | 128.55 | 9.54 | 0.27 | 0.01 | 0.59 |

**Table 2B.** Impact of donor and supplementation on bacterial community metrics, following a short-term incubation (16 h) with either celecoxib (CX) or carrier (PEG), n = 8.

| Metric | Treatment |  | SEM | *P* value |  |  |
| --- | --- | --- | --- | --- | --- | --- |
|  | CX | PEG |  | Treatment | Donor | Treatment*Donor interaction |
| Evenness | 0.55 | 0.59 | 0.01 | 0.06 | 0.003 | 0.38 |
| Shannon ($H'$) | 2.54 | 2.70 | 0.06 | 0.08 | 0.001 | 0.33 |
| Simpson | 0.84 | 0.86 | 0.009 | 0.14 | 0.009 | 0.10 |
| Inverse Simpson ($D_2$) | 7.59 | 8.13 | 0.63 | 0.55 | 0.02 | 0.23 |
| Fisher's diversity | 13.28 | 13.17 | 0.25 | 0.49 | <0.0001 | 0.34 |
| Total Richness (S) | 92.56 | 97.12 | 2.78 | 0.88 | 0.005 | 0.07 |
| Chao | 94.01 | 114.65 | 5.34 | 0.01 | 0.008 | 0.89 |

### Short-term exposure to celecoxib increases number of community metabolic networks

Association networks were constructed to highlight potential shifts among bacterial relative abundances after short-term exposure to CX or its vehicle. The number of links in the network was lower when the vehicle PEG alone was provided (Figure 4A), but the central position of *Phascolarctobacterium*, *Parabacteroides*, *Coprococcus* 2 and Unclassified Victivallales and the linkages among these genera suggest concomitant changes in their relative abundances in presence of PEG. The central position of the acetate-producing *Erysipelatoclostridium* in the celecoxib network (Figure 4B) indicated that its relative abundance was associated with *Subdoligranulum*, a butyrate and lactate-producing genus, in a donor-dependent manner (*P* < 0.05, Supplementary Table 4).

**Figure 4.**
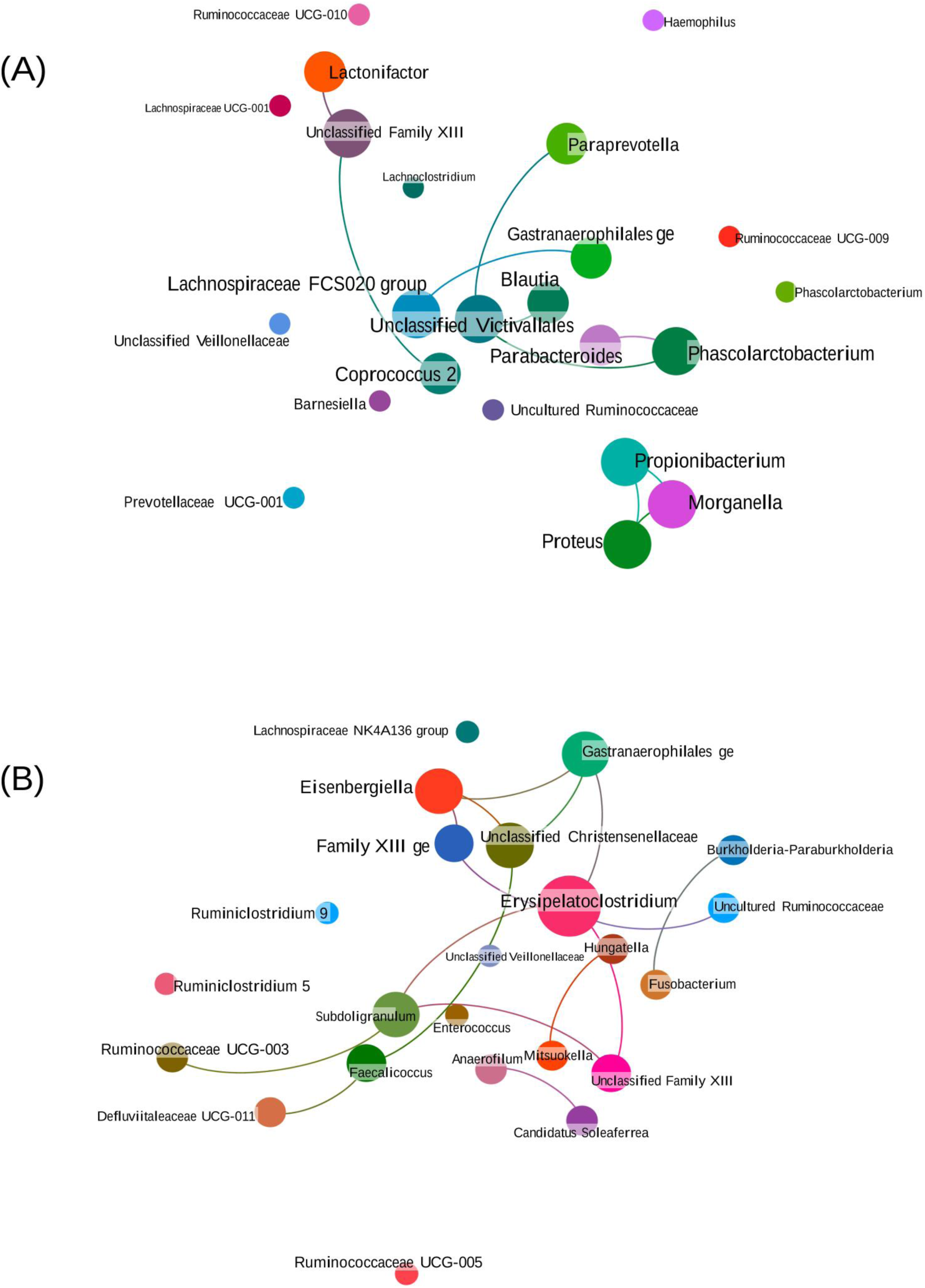
Bacterial network of communities exposed to short-term administration of celecoxib (CX) or its vehicle (control treatment, PEG). (A) The faecal microbiota exposed only to PEG displayed a lower number of associations, while the network architecture was enriched when CX was provided (B). This representation uncovers relationships between clusters of genera across donors. Nodes representing bacterial genera with similar abundances tended to cluster closely. Node diameter indicates the relative abundance of each genus. Neighbourhood selection method (MB) was the graphical model inference procedure selected. The associations described may be considered ‘core’ functional relationships preserved across donors.

Networks connecting bacterial load, metabolites and relative abundances were assembled to monitor functional alterations associated with changes in genera abundances. Activity-based relative abundance of *Bacteroides* was positively associated with butyrate, propionate and valerate levels and total 16S rRNA gene copy number in presence of PEG (Figure 5A, Supplementary Table 4). These results confirmed that *Bacteroides* was one of the most abundant active genera, as first suggested in Figure 3. Higher number of links in comparison with PEG were detected after 16 h of exposure to celecoxib (Figure 5). Indeed, one network linking acetate with the relative abundance of diverse taxa was only present when celecoxib was supplied and contained the largest number of active genera participating. Relative abundance of *Faecalibacterium* decreased over time with celecoxib supplementation, corresponding with the decline in butyrate levels (Figure 3). However, this genus tended to be positively associated with acetate production (Figure 5B). Relative abundance of active *Lachnoclostridium* was significantly associated with butyrate in the celecoxib treatment (Figure 5B). Here, *Oscillospira*, *Allisonella*, *Hungatella* and *Streptococcus* were linked because they possibly participated in butyrate production as well. *Megamonas* and *Megasphaera* were other genera with high relative abundances positively associated with branched-chain SCFA and isovalerate. The associations between the latter metabolites and *Colinsella* remained unchanged upon celecoxib exposure, suggesting that this community function is either independent from drug supplementation or directly associated with PEG addition (Figure 5A and 5B). Similarly, associations between *Allisonella* (butyrate consumer), *Streptococcus* (butyrate producer) and butyrate were common for both treatments. Results from the network analysis indicate that shifts in the community metabolic activity upon short-term supplementation of celecoxib, potentially impacted butyrate production.

**Figure 5:**
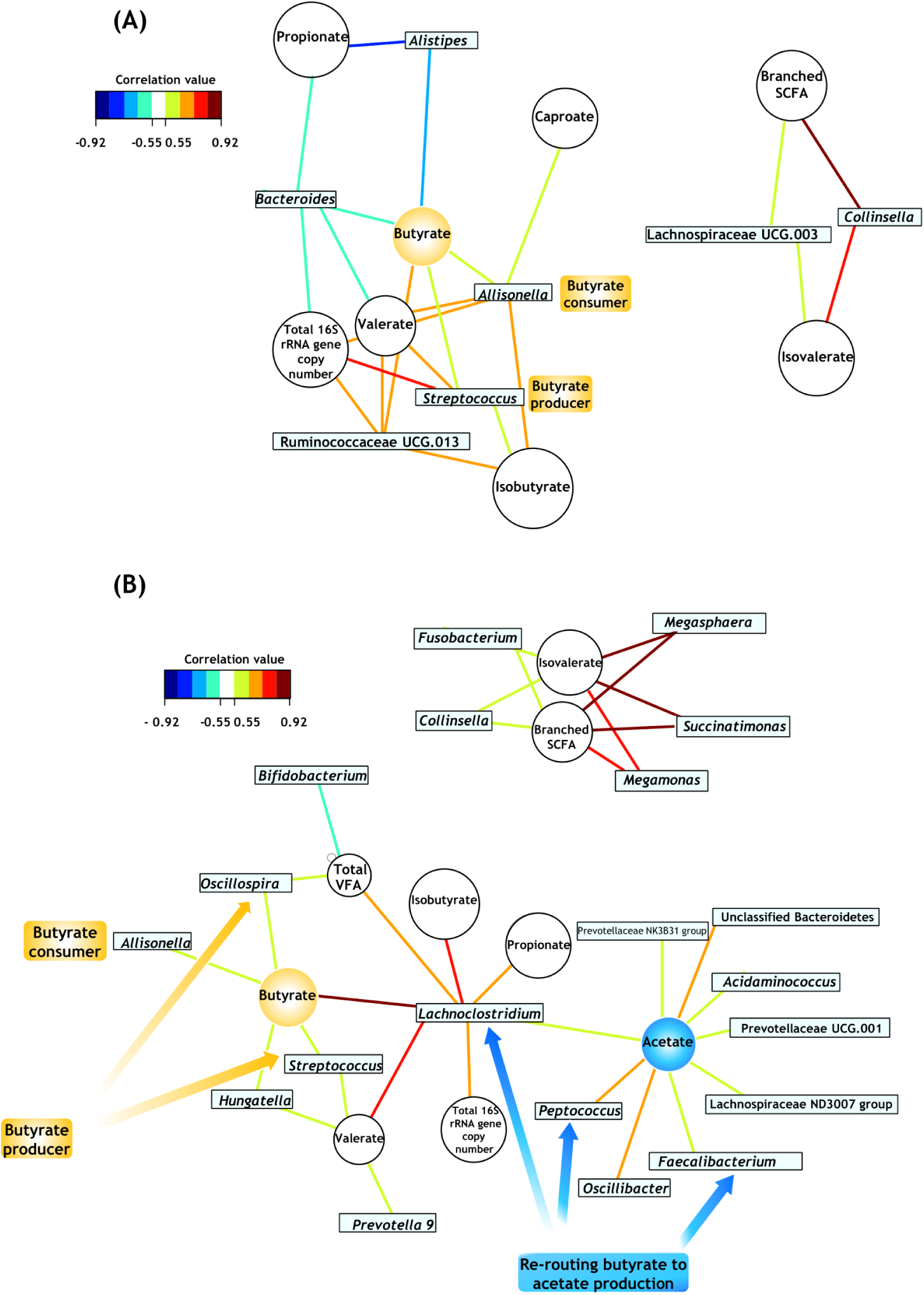
Community metabolic activity indicates that networks of bacterial interactions were few when only the vehicle (PEG) was supplied (A) and increased following short-term supplementation of celecoxib (B). These bipartite networks are based on the regularised canonical correlations between relative bacterial abundances and relative concentrations of the main short-chain fatty acids. Interactions have been filtered for an absolute correlation above 0.5 and are coloured following the key shown. Significant interactions are indicated by shorter lines, and genera with similar abundances within treatment tended to cluster closely.

### Inter-individual differences play a role in reduction of *in vitro* inflammation when celecoxib is presented to the microbiome

Understanding the microbial interactions with the host epithelium is essential to assess whether the anti-inflammatory efficacy of celecoxib remains unaltered following exposure to the simulated microbiome. We incorporated the immune component in a simplified inflammation model using THP-1 (macrophage-like) cells. We further verified the microbiome impact using a multicellular model simulating the enterohepatic ecosystem, including enterocyte-like and goblet-like cell lines, plus THP-1 cells. This set-up enabled closer communication between cell lines, as would occur when macrophages infiltrate the underlying intestinal mucosa.

Direct contact of macrophage-like cells with filter-sterilized supernatants from the CX-exposed microbiota resulted in a variable outcome in IL-8 and CXCL16 concentrations. Compared with the PEG controls, a decrease in IL-8 and CXCL16 was only observed in 4 out of 8 individuals (Figure 6A and 6B).

**Figure 6.**
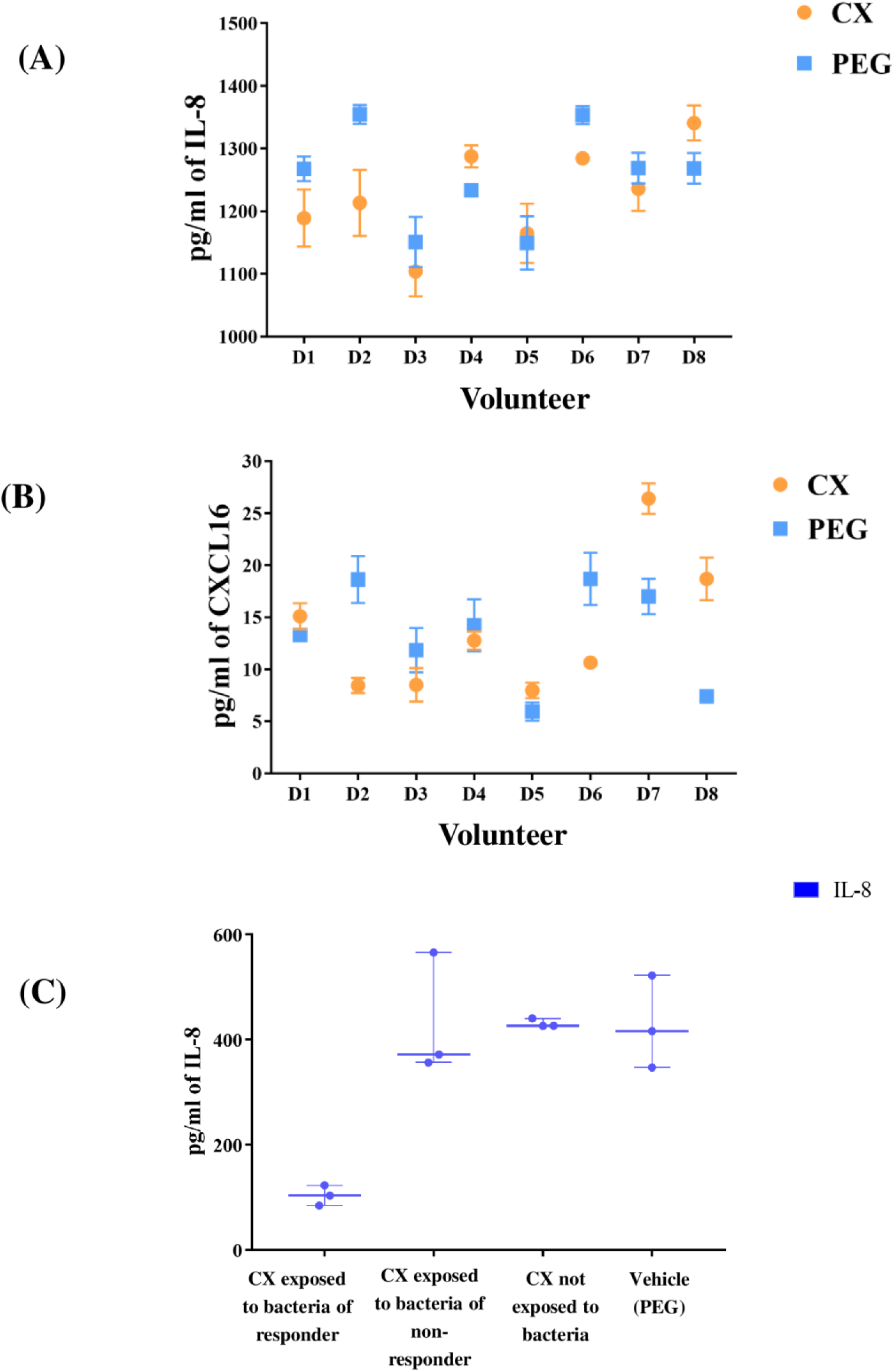
Concentration of pro-inflammatory cytokines was decreased in a donor-dependent manner when celecoxib was supplied. Volunteers are numbered consecutively as D1-D8. (A) IL-8 was significantly lower in the simplified inflammation model subjected to the bacteria-exposed celecoxib from 3 of the volunteers screened (D1, D2 and D6). Two additional volunteers (D3 and D7) showed a trend for decreased IL-8, but it was not significantly lower than the control treatment. (B) CXCL16 was significantly lower in the simplified inflammation model subjected to the bacteria-exposed celecoxib from only two volunteers (D2 and D6) and tended to be decreased in other two (D3 and D4). (C) The two volunteers with the most extreme metabolic responses in butyrate production were selected to be screened in a multicompartment cell model resembling the enterohepatic system. Volunteer 6 (D6) had the most significant decrease in butyrate levels and was considered “responder” to the CX treatment. Volunteer 8 (D8) did not show any significant difference in butyrate between the control and the treatment after 16 h of incubation and it was considered as “non-responder”. IL-8 concentration was significantly lower in the model subjected to the supernatant of D6 containing bacteria-exposed celecoxib (“responder volunteer”). The opposite effect was observed in D8 (“non-responder volunteer”), suggesting that interindividual differences in the functionality of the microbiome may play a role in the response to anti-inflammatory drugs.

High IL-8 concentrations were found when non-bacterially exposed celecoxib was applied in the enterohepatic model (Figure 6C). In contrast, the lowest concentration of IL-8 and thus the most significant decrease in inflammatory response was found with the celecoxib-exposed supernatant of donor 6 (*P* < 0.05, Table 3). Celecoxib that was not in contact with bacteria promoted the highest decrease in transepithelial electrical resistance (TEER) (*P* < 0.05, Table 3), in comparison with supernatants from celecoxib-exposed bacteria. This outcome indicates less damage to epithelial barrier function and decreased inflammatory response when gut microbiota is exposed to celecoxib.

**Table 3.**
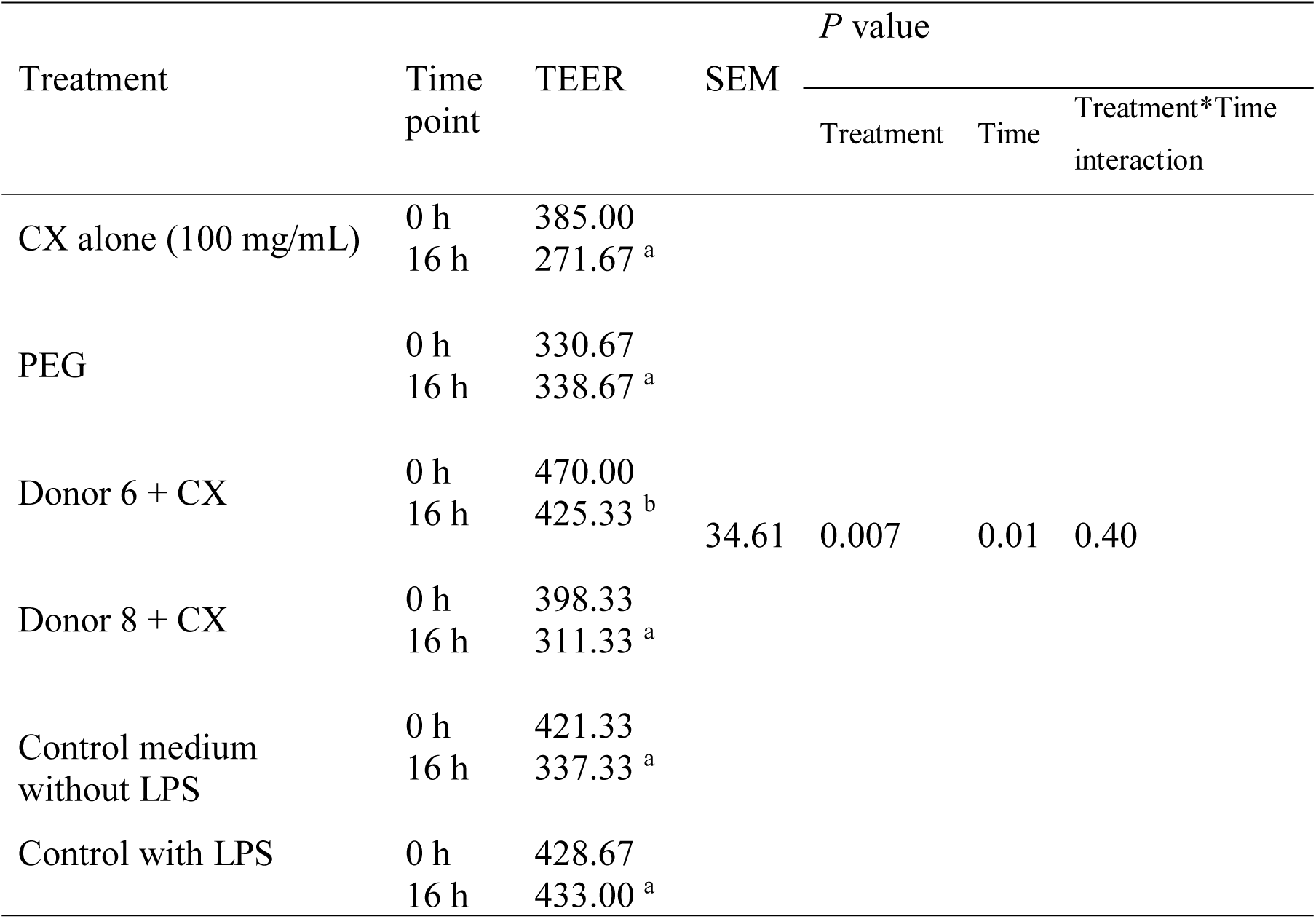
Epithelial integrity metrics of a multicompartment cell model simulating the enterohepatic metabolism, after 16 h of incubation with CX-exposed bacteria. 0 h = prior to LPS exposure, 16 h = after 16 h of incubation with LPS. Different superscripts indicate significantly different means across treatments, within a particular time point. N = 3.

## DISCUSSION

As the lipophilic characteristics of celecoxib favour metabolic elimination (19), exploring only the pathways for celecoxib distribution in the host is insufficient for assessing drug bioavailability. *In vitro* models have been applied to simulate drug-bug associations in the human gut (20, 21), but the impact of NSAIDs on the metabolic activities of the microbiome has been overlooked. Previous studies reported the influence of different NSAIDs on the gut microbiome (22), yet no changes in bacterial richness, microbiome composition or beta diversity were found when celecoxib was supplied (16). These results focused on screening community structure using DNA-based amplicon sequencing, which may also include DNA of dead bacterial cells. As such approach may deliver a partial overview, we used cDNA as an indicator of the metabolically active bacterial population and we performed community composition analyses based on information from transcribed RNA (23). In this way, we intended to highlight the main bacterial genera whose metabolic activities were impacted by celecoxib in the RNA pool. Activity-based cell load was not significantly different, and the only alterations were on the bacterial metabolic activities. Although composition, diversity and richness were not significantly different as result of the treatment, differences in the most abundant genera between PEG and celecoxib were observed. PERMANOVA analysis suggested that donor was the main driver for dissimilarities in community structure.

The shifts in microbial metabolism observed in our study may indicate that celecoxib can inhibit functional activities of bacterial groups, rather than of single genera, ultimately impacting microbial fermentation (24). This was confirmed by the total active bacterial load, which remained unchanged between treatments. Celecoxib users have been reported to have enrichment of Enterobacteriaceae and Acidaminococcaceae (22). Here, we validated that the relative abundance of transcriptionally active *Escherichia*-*Shigella* (Enterobacteriaceae) was increased with celecoxib supplementation, whereas the relative abundance of active *Acidaminococcus* (Acidaminococcaceae) was positively associated with acetate in the same treatment. These results highlight the applicability of our *in vitro* model used for this study. Coriobacteriaceae (*Colinsella*) has been proposed to participate in xenobiotic metabolism by decreasing levels of serine and glycine (24), which can be both fermented to acetate in the gut (25). We observed that the relative abundance of this genus was positively associated with isovalerate levels when celecoxib was added. As indicated in the corresponding KEGG module, *Colinsella aerofaciens* has the capacity of using isovalerate for synthesising leucin and serine (26), confirming its potential xenobiotic role. On the other hand, the capacity of *Alistipes* for producing succinate (27), may explain its association with propionate in the PEG treatment. *Dialister* is another succinate producer (28) present at high relative abundance, but it does not seem to participate in the propionate network when PEG was supplied. This may suggest that, even based on cDNA, high relative abundance may not be an indicator of metabolic activity but just of maintenance metabolism.

The significantly lower butyrate concentration post-celecoxib delivery may have several implications. Butyrate has tumour inhibitory properties (29) but its paradoxical correlation with polyp formation (30, 31) suggests that the CRC-butyrate interplay is not just a direct effect of altered SCFAs concentrations, and may also be associated with changes in microbiome functionality. Indeed, the protecting properties of celecoxib towards CRC (32, 33) may be partially derived from lowering bacteria-derived butyrate. Nutrients can be utilized in their final form by both host and gut microbes, leading to competition for resources (34). In fat, blocking fatty acid supply and host recapture has been proposed as strategy for influencing cancer cell bioenergetics (35, 36), but few studies have addressed the potential role of acetate in cancer (37). Acetyl-CoA is the main carbon source in mammalian cells, and the ligation of acetate and CoA by acetyl-CoA synthetase (ACSS) contributes to the supply of this metabolite (37). Cancer cells use acetyl Co-A during aerobic metabolism regardless of stress conditions (37). Butyrate is also metabolised to acetyl-CoA (29), inhibiting tumour growth via silencing acetyl-CoA synthetases (35) or potentially through acetate uptake. As acetyl-CoA is a bacterial substrate for fatty acid production (38), our network analysis suggests that acetate was used for bacterial metabolic functions when celecoxib was supplied. Thus, celecoxib may favour bacterial competition for acetyl Co-A, generating potential host protective effects. Indeed, microbiota-derived acetate has been associated with inhibition of cancer-associated deacetylation (39). Increasing acetate concentration shifts SCFA ratios (40), decreasing butyrate and explaining why the acetate:propionate:butyrate ratios tended to change when celecoxib was supplied. *Faecalibacterium* sp. is a well-known butyrate producer and acetate consumer (40) and its presence in the acetate network may indicate that the relative abundance of this genus increased when celecoxib was provided and acetate levels increased.

Synthesis of reduced products including H_2_, succinate, and butyrate creates redox balance during colonic fermentation, whereas formation of oxidised products, such as acetate, is associated with bacterial ATP production (38). Fermentative activities of *Lachnoclostridium* yield ethanol, acetate, CO_2_ and H_2_ (41, 42), explaining the numerous links with other genera in the celecoxib network. These potential cross-feeding interactions (43, 44), may support its positive association with the total 16s rRNA gene copy number. Acetate is a co-substrate for butyrate production (45), but butyrate production may be reduced when NADH used for H_2_ formation during glycolysis is not available to convert acetoacetyl-CoA to butyryl-CoA (38). Acetate-producing *Lachnoclostridium* and *Oscillospira* may have been involved in the shift from butyrate to acetate production when celecoxib was supplied. This is because these genera possess transporters involved in uptake of oligosaccharides and glycerol-P (43). Additionally, Peptococcaceae uses the acetyl-CoA pathway in a reverse direction to oxidize butyrate (46). Acetyl Co-A is consequently routed into acetate formation (45), which is energetically advantageous for bacterial use (38). In fact, butyrate producers may shift metabolic pathways towards acetate under stress conditions (47). As in our study, other acetate producers such as Bacteroidetes and Prevotellaceae positively correlated to butyrate-producing communities using the acetyl CoA pathway for butyrate production (46), supporting the observed shift in butyrate towards acetate (48).

Although the role of SCFA on inflammation and CRC progression has been documented, most of the studies lack natural commensal microbiota cross-talking with host cells (29, 31). As interindividual differences in microbiome play a crucial role on drug responses (12), elucidating whether microbiome fermentation products synergistically act with NSAIDs for impacting inflammation is paramount. Celecoxib was responsible for only 5% of the variance in bacterial community composition among the 8 donors, but the effect of celecoxib-exposed bacteria on the proinflammatory IL-8 was sufficient to be detected in a donor-dependent manner. Moreover, samples derived from 50% of the donors showed increased CXCL-16, which is a marker of favourable prognosis in CRC (49). This suggests that the interaction between celecoxib and microbiome to resolve inflammation may be not solely related to community composition, and functionality must be accounted for. While butyrate seemed to be reduced with short-term treatment, Acetate:Propionate:Butyrate ratio and relative levels of acetate (of the total SCFA) may be other functional bacterial metrics to be considered for assessing successful reduction on inflammation when NSAIDs are administered.

As indicated in our study, gut microbiome activities and its interplay with the host can be modulated through celecoxib. Our *in vitro* model retained interindividual variability, allowing for mechanistic research on drug-microbiota-host interplay, fundamental for developing individualised therapeutic strategies. Divergence in the molecular approaches (i.e. next generation sequencing, qPCR) used for investigating CRC-associated microbiome signatures contributes to the lack of consensus on the composition of microbiota in CRC patients. Information on the role of SCFA in cancer signalling remains inconclusive, and thus, future studies assessing the crosstalk of the microbiota and colon cells in CRC are imperative. Certainly, individual microbiome may need to be considered before supplementing celecoxib as a colon targeted drug for CRC. However, focusing on functionality rather than taxonomic diversity may be the next frontier for developing predictive markers of CRC-induced dysbiosis and NSAID efficacy.

## MATERIALS AND METHODS

### Faecal incubations simulating the proximal colon environment

Batch incubations of faecal samples were conducted using phosphate-buffered nutritional medium (pH 5.9, 0.1 M) containing (per litre): 12.25 g of KH_2_PO_4_ (Carl Roth GmbH, Karlsruhe, Germany), 1.78 g of Na_2_HPO_4_ (Carl Roth GmbH, Karlsruhe, Germany), 3 g of yeast extract (Oxoid Ltd, Basingstoke, Hampshire, England), 1 g of proteose peptone (Oxoid Ltd, Basingstoke, Hampshire, England), 0.25 g of gum Arabic from acacia tree (Sigma Aldrich Co., St-Louis, USA), 0.5 g of apple pectin (Sigma Aldrich Co., St-Louis, USA), 0.25 g of xylan (Carl Roth GmbH, Karlsruhe, Germany), 1 g of potato starch (Prodigest, Zwijnaarde, Belgium), and 4 g of mucin from porcine stomach Type II (Sigma Aldrich Co., St-Louis, USA).

Faecal material was obtained from eight healthy, non-obese, non-smoking volunteers (4M and 4F, 30 ± 2.8 yo), following an omnivorous diet and without antibiotic exposure for the prior 6 months. Stool samples were collected and prepared as previously described (50). Celecoxib (Janssen Pharmaceutica, Beerse, Belgium) was dissolved in 70:30 (v/v) PEG_400_:H_2_O as vehicle, at a stock concentration of 100 mg/mL. For each donor, triplicate incubations of 10% faecal slurry, 100 mg/mL of celecoxib or 70:30 (v/v) PEG_400_ (Acros, Geel, Belgium):H_2_O, and nutritional medium (final volume = 175 mL) were performed for 16 hours at 37°C and 120 rpm (New Brunswick Scientific InnOva 4080 Incubator Shaker, Wezembeek-Oppem, Belgium), under anaerobic conditions (90%-10% N_2_/CO_2_). Once the incubation was completed, samples were collected, flash-frozen using liquid nitrogen and preserved at –80 °C for further analysis.

### Surveying community functionality and composition

#### Short-chain Fatty Acids (SCFAs) production

SCFA were used as benchmarks of community activity. Extraction and analyses have been described by Truchado et al. (51). Differences in SCFA concentrations among treatments were compared using a repeated measures mixed model (52), with the lsmeans adjustment and Bonferroni correction for multiple comparisons (SAS, 2012). Statistical significance was assumed at *P* < 0.05.

#### Quantification of total metabolically active bacterial community

Samples collected from the faecal incubations at 0h (start) and after 16h (end of experiment) were thawed on ice, centrifuged for 5 minutes at 4°C and 5000 × *g*, and disrupted two times for 25 seconds at 1400 rpm (Power Lyser 24, MO BIO Laboratories, Carlsbad, USA), with a 5-second interval, where the samples were placed on ice. RNA was further extracted using the Nucleospin RNA plus kit (Macherey Nagel, Düren, Germany), following the manufacturer’s instructions. Reverse transcription was completed using the Reverse Transcriptase Core kit (Eurogentec, Seraing, Belgium).

For the quantification of total bacterial 16S rRNA gene copy numbers, a standard curve was constructed using serial dilutions of plasmid DNA from a clone identified as *Bifidobacterium breve* LMG 33264. Briefly, universal bacterial primers 27F and 1492R were used to amplify the full-length 16S rRNA gene from the plasmid DNA of a *Bifidobacterium breve* clone. The resultant PCR product was purified using the using the InnuPREP PCRpure Kit (AJ innuscreen, Berlin, Germany), cloned in *E. coli* with the TA Cloning Kit (Life Technologies, Carlsbad, VS) and plasmid extracted with the PureYield Plasmid Miniprep System (Promega, Leiden, The Netherlands). The mass concentration of the plasmid was measured using an ND-1000 spectrophotometer (NanoDrop Technologies, Wilmington, DE), converted to the molecule concentration (53) and copy numbers of total bacteria in 50 ng of cDNA and per ml of faecal incubation were determined by relating the threshold cycle (*C_T_*) values to the standard curves based on the following equation: *Y* = −3.193 × log*X* + 35.003 (Y, *C_T_* value; *X*, copy number of 16S rRNA gene) (*r*^2^ = 0.996) (53).

Quantitative reverse transcription-PCR (qRT-PCR) analysis of the metabolically active community was achieved with the StepOnePlus Real-Time PCR system (Applied Biosystems, Foster City, CA USA). Amplification reactions were in: 10× Taq buffer with KCl, 25 mM of MgCl_2_, 10 mM of dNTP mix, 10 µM of each forward and reverse primers, 20 mg/mL of BSA (Thermo Scientific, Waltham, USA), 1.25 U of Taq Polymerase, 50 ng of cDNA and 0.1× SYBR Green I ([Invitrogen, Carlsbad, USA] provided at 10 000×, stock solutions of 20× were prepared in DMSO). qRT-PCR conditions included 10 minutes at 95°C, followed by 40 cycles of 15 s at 95°C and 30 s at 60°C. The efficiencies (*E*) of RT-PCR were calculated from the given slopes in StepOnePlus software using the following equation: *E* = [10(^−1/slope^) − 1] × 100%. Data generated from reactions with efficiencies between 90 and 110% were used for further analysis (53).

### Community composition and dynamics

The V5-V6 hypervariable region of the 16S rRNA gene from the cDNA was amplified using primers 341F and 785R. High-throughput amplicon sequencing was performed with the Illumina MiSeq platform according to the manufacturer’s guidelines at LGC Genomics GmbH (Berlin, Germany). Library preparation and purification as well as bioinformatic processing are described in the supplemental material.

Data was imported into R using *phyloseq* (54) and taxon abundances were rescaled by calculating the taxon proportions and multiplying them by the minimum sample size (n = 2201) present in the data set (55). Inverse Simpson was the metric used for assessing alpha diversity and Pielou’s index was used as indicator of community evenness (56). Differences in alpha diversity and evenness measures were compared among treatments using a repeated measures mixed model in SAS (version 9.4, SAS Institute, Cary, USA). Beta diversity based on Chao and Bray-Curtis indices was used to examine dissimilarity and determine the impact of treatment and time on microbial community structure (52). PCoA was employed to visualize the differences among samples, using the vegan package in R (57). Stratified permutational multivariate analysis of variance (PERMANOVA) with 999 permutations was conducted to indicate the significance of each covariate (time or treatment) on the microbial community.

ANOVA was applied to reveal whether the distribution of the genera was different between treatments over time (57). Because of the over-dispersion in the OTU data, a zero-inflated negative binomial distribution count model was used to assess the effect of donor, time and treatment and the interactions between donor*time*treatment on each individual genus. The model was selected based on the Akaike Information Criterion (AIC). Differences among library size sample were accounted for with the offset option in proc GLIMMIX in SAS (52).

Sparse inverse covariance selection was employed to infer an association network within treatments, using the package SpiecEasi (v0.1.2) (58) in R (v3.5.1). Neighbourhood selection method (S-E(MB) was used to construct a network of bacterial genera displaying core networks with either functional or phylogenetic similarities. The calculated adjacency matrix was examined with the software Gephi 0.9.2 (59), using the Fruchterman-Reingold algorithm for visualizing significant correlations (R ≥ 0.7). This method computes a layout in which the length of the connections indicates the absolute value of the correlation. The nodes representing genera whose relative bacterial abundances have several significant correlations among them were placed closer to each other in the network (50).

Bipartite networks highlighted functional associations among bacterial genera and metabolites (52), using a pair-wise similarity matrix obtained from a Regularised Canonical Correlation Analysis (60). Values of the similarity matrix were computed as the correlation between relative abundances of bacterial genera and metabolic variables, projected onto the space spanned by the first 3 components retained in the analysis (r ≥ 0.75). Genera in the plot were close to correlated variables in the treatment where they were more abundant (52).

### Assessing the anti-inflammatory potential of bacteria-exposed celecoxib

#### Simplified inflammation model

The human monocytic leukaemia cell line THP-1 (THP-1, ECACC 88081201, Public Health England, UK) was grown in Roswell Park Memorial Institute (RPMI) 1640 culture medium (Thermo Fisher Scientific, Waltham, USA) supplemented with 10% heat-inactivated foetal bovine serum (iFBS) (Greiner Bio-One, Wemmel, Belgium) and 1% antibiotic-antimycotic solution (10,000 units/mL of penicillin, 10,000 µg/mL of streptomycin, and 25 µg/mL of Amphotericin B) (Thermo Fisher Scientific, Waltham, USA), at 37°C in 10 % CO_2_ in a humidified incubator. THP-1 cells were sub-cultured twice a week and used between passage 20 and 30.

Macrophage-like phenotype was obtained by treating cells with 25 ng/mL phorbol 12-myristate 13-acetate (PMA, Sigma Aldrich, St-Louis, USA) for 48 h in 24-wells culture plates (Corning, Sigma Aldrich, St-Louis, USA) at a density of 1 x 10^5^ cells/cm^2^. Differentiated, plastic-adherent cells were washed twice with culture medium (supplemented RPMI 1640 without PMA) and incubated for 24 h. Medium was removed and replaced by supernatant from the eight incubations, passed through a sterile 0.2 µm filter and diluted five times with RPMI 1640 without supplementation. Celecoxib solution at 100 mg/mL dissolved in 70:30 (v/v) PEG:H_2_O or 70:30 (v/v) PEG:H_2_O alone were diluted 1:5 with medium and added as controls of non-bacteria exposed celecoxib and vehicle. This dilution step was performed to simulate low solubility and the small aqueous volume in the colon (14). LPS (100 ng/mL) dissolved in RPMI 1640 was added to the cells to simulate systemic inflammation, as in CRC. The control used in all measurements was PMA-differentiated THP-1 macrophages exposed to the culture medium.

#### Development of an *in vitro* model of the enterohepatic system

A novel model including intestinal epithelial (Caco-2, ECACC 86010202), goblet (HT29-MTX-E12 ECACC 12040401, Public Health England, UK), hepatic (HepG2 ATCC HB-8065, Molsheim, France), and immune-like cells (THP-1, ECACC 88081201, Public Health England, UK) was developed to study host-drug-microbiome interactions (61, 62). Cell layers were grown in a double-chamber insert (Corning, Sigma Aldrich, St-Louis, USA), and maintained for 23 days. Briefly, 100 µl of a THP-1 cell suspension (10^6^ cells/mL) in 20 mg/mL collagen from rat tail type I (Sigma Aldrich, St. Louis, USA) were spread in the basolateral surface of inverted inserts. After 45 min of incubation at 37°C and 10% CO_2_, a 9:1 cell suspension of Caco-2/HT29-MTX cells (density of 7.5×10^4^ cells/cm^2^) was added to the apical side of the insert in 1.5 ml of Dulbecco’s Modified Eagle’s medium (DMEM) supplemented with 10% (v/v) of iFBS and 1% of antibiotic-antimycotic solution (v/v). The basolateral compartment of the double-chamber system was filled with 2 ml of supplemented RPMI 1640, and the medium was refreshed every 48 h. Inserts were transferred to 6-well plates with pre-seeded HepG2 cells at day 19 post-seeding, when the cell culture media in the basal compartment was replaced by RPMI 1640, and the co-culture was maintained 24 h prior to the assay. HepG2 cells were seeded at a density of 10^5^ cells/cm^2^ and maintained 5 days in supplemented minimal essential medium (MEM) supplemented with 10% (v/v) of iFBS and 1% (v/v) of antibiotic-antimycotic solution. Five days prior the assay all the cell culture media was depleted of antibiotic/antimycotic solution. Supernatants from volunteers (2F, 29 yo) showing extreme microbiota metabolism responses in the short-term incubations were used for validating whether celecoxib effectively decreases inflammation even after exposure to the gut microbiota. Volunteer 6 displayed the most significant decrease in butyrate after 16 h of incubation, while volunteer 8 did not present any changes. Triplicate supernatants from each volunteer were passed through a sterile 0.2 µm filter, diluted five times with DMEM without supplementation, pH was adjusted to pH 7-7.2 with filter-sterilized NaOH 0.5M (Carl Roth GmbH & Co), and placed in the apical compartment of the model. Results from the simplified inflammation model indicated that there were no significant differences between control and treatment among donors. Thus, we only included celecoxib solution dissolved in 70:30 (v/v) PEG:H_2_O or 70:30 (v/v) PEG:H_2_O alone in the enterohepatic model. Celecoxib solution at 100 mg/mL and a 70:30 (v/v) PEG:H_2_O were diluted 1:5 with DMEM without supplementation, and added to the apical compartment, as controls of non-bacterially modified celecoxib and vehicle. This dilution step was performed to simulate low solubility and the small aqueous volume in the colon (14). LPS (100 ng/mL) was added to the basal compartment to simulate systemic inflammation. Plates were incubated for 16 h at 37°C, 10% CO_2_ and 95% humidity. TEER was evaluated with the Millicell ERS-2 Voltohmmeter at the beginning and at the end of the assay (Merck-Millipore, Overijse, Belgium).

#### Evaluating inflammatory response

Concentration of IL-8 and CXCL16 were measured in the simplified inflammation model with the Human IL-8 (CXCL-8) and Human CXCL16 Mini ABTS ELISA Development Tests (PeproTech, London, UK) and the ABTS ELISA Buffer Kit (PeproTech, London, UK), following the manufacturer’s instructions. Samples were not diluted, and determination was performed at 405 nm, with correction for 650 nm (Infinite 200 PRO, Tecan, Männedorf, Switzerland). IL-8 was additionally determined in the presystemic metabolism model, using samples from the basal compartment diluted ten times with PBS (ABTS ELISA Buffer Kit, PeproTech, London, UK), and measured as indicated above.

### Data Availability

Raw 16S rRNA reads have been made available on the SRA under accession number ID PRJNA540406. Original R scripts are available in GitHub (https://github.ugent.be/ehernand/CX_sequencing).

## ACKNOWLEDGEMENTS

E.H.-S. is a postdoctoral fellow supported by Flanders Innovation and Entrepreneurship (Agentschap Innoveren & Ondernemen). M.C. was supported by an FWO postdoctoral fellowship (FWO/12R2717N). E.H.-S. and T.Vd.W. conceived the simulations of the colon environment. E.H.-S. and M.C. devised cell models. J.S. and L.L. supplied material and advised on the use of celecoxib. E.H.-S., E.H., and M.C. conducted all laboratory work. R.P. completed the amplicon sequencing library pre-processing and guided sequencing data mining. E.H.-S. performed the statistical analyses and interpretation of all data generated, produced figures and tables, and wrote the manuscript. All authors read and approved the manuscript. The authors gratefully acknowledge the volunteers participating in this study. We thank Dr. Davide Gottardi for his scientific advice and Racha El Hage, Fabian Mermans, Olivier Dezutter, and Dr. Chiara Ilgrande for their technical assistance. The authors declare that they have no competing interests.

## REFERENCES

1. Kumar M, Cash BD. 2017. Screening and surveillance of colorectal cancer using CT colonography. Current treatment options in gastroenterology 15:168–183.

2. Arnold M, Sierra MS, Laversanne M, Soerjomataram I, Jemal A, Bray F. 2017. Global patterns and trends in colorectal cancer incidence and mortality. Gut 66:683–691.

3. Wang J, Cho NL, Zauber AG, Hsu M, Dawson D, Srivastava A, Mitchell-Richards KA, Markowitz SD, Bertagnolli MM. 2018. Chemopreventive efficacy of the cyclooxygenase-2 (Cox-2) inhibitor, celecoxib, is predicted by adenoma expression of Cox-2 and 15-PGDH. Cancer Epidemiology and Prevention Biomarkers 27:728–736.

4. Bhala N, Emberson J, Merhi A, Abramson S, Arber N, Baron J, Bombardier C, Cannon C, Farkouh M, FitzGerald G. 2013. Vascular and upper gastrointestinal effects of non-steroidal anti-inflammatory drugs: meta-analyses of individual participant data from randomised trials. Elsevier.

5. Kawanishi S, Ohnishi S, Ma N, Hiraku Y, Murata M. 2017. Crosstalk between DNA damage and inflammation in the multiple steps of carcinogenesis. International journal of molecular sciences 18:1808.

6. Wang D, DuBois RN. 2010. The role of COX-2 in intestinal inflammation and colorectal cancer. Oncogene 29:781.

7. Funk CD, FitzGerald GA. 2007. COX-2 inhibitors and cardiovascular risk. Journal of cardiovascular pharmacology 50:470–479.

8. Cuzick J, Otto F, Baron JA, Brown PH, Burn J, Greenwald P, Jankowski J, La Vecchia C, Meyskens F, Senn HJ. 2009. Aspirin and non-steroidal anti-inflammatory drugs for cancer prevention: an international consensus statement. The lancet oncology 10:501–507.

9. Johnson CH, Spilker ME, Goetz L, Peterson SN, Siuzdak G. 2016. Metabolite and microbiome interplay in cancer immunotherapy. Cancer research 76:6146–6152.

10. Tjalsma H, Boleij A, Marchesi JR, Dutilh BE. 2012. A bacterial driver–passenger model for colorectal cancer: beyond the usual suspects. Nature Reviews Microbiology 10:575.

11. Spanogiannopoulos P, Bess EN, Carmody RN, Turnbaugh PJ. 2016. The microbial pharmacists within us: a metagenomic view of xenobiotic metabolism. Nature Reviews Microbiology 14:273.

12. Wilson ID, Nicholson JK. 2017. Gut microbiome interactions with drug metabolism, efficacy, and toxicity. Translational Research 179:204–222.

13. Sousa T, Paterson R, Moore V, Carlsson A, Abrahamsson B, Basit AW. 2008. The gastrointestinal microbiota as a site for the biotransformation of drugs. International journal of pharmaceutics 363:1–25.

14. Philip AK, Philip B. 2010. Colon targeted drug delivery systems: a review on primary and novel approaches. Oman medical journal 25:79.

15. Srisailam K, Veeresham C. 2010. Biotransformation of celecoxib using microbial cultures. Applied biochemistry and biotechnology 160:2075–2089.

16. Bokulich NA, Battaglia T, Aleman JO, Walker J, Blaser MJ, Holt PR. 2016. Celecoxib does not alter intestinal microbiome in a longitudinal diet-controlled study. Clinical Microbiology and Infection 22:464–465.

17. Hardy JG, Wilson CG, Wood E. 1985. Drug delivery to the proximal colon. Journal of Pharmacy and Pharmacology 37:874–877.

18. Macfarlane G, Macfarlane S, Gibson G. 1998. Validation of a three-stage compound continuous culture system for investigating the effect of retention time on the ecology and metabolism of bacteria in the human colon. Microbial ecology 35:180–187.

19. Paulson SK, Vaughn MB, Jessen SM, Lawal Y, Gresk CJ, Yan B, Maziasz TJ, Cook CS, Karim A. 2001. Pharmacokinetics of celecoxib after oral administration in dogs and humans: effect of food and site of absorption. Journal of Pharmacology and Experimental Therapeutics 297:638–645.

20. McConnell EL, Fadda HM, Basit AW. 2008. Gut instincts: explorations in intestinal physiology and drug delivery. International journal of pharmaceutics 364:213–226.

21. Esch EW, Bahinski A, Huh D. 2015. Organs-on-chips at the frontiers of drug discovery. Nature reviews Drug discovery 14:248.

22. Rogers MA, Aronoff DM. 2016. The influence of non-steroidal anti-inflammatory drugs on the gut microbiome. Clinical Microbiology and Infection 22:178. e1–178. e9.

23. Blazewicz SJ, Barnard RL, Daly RA, Firestone MK. 2013. Evaluating rRNA as an indicator of microbial activity in environmental communities: limitations and uses. The ISME journal 7:2061.

24. Montrose DC, Zhou XK, McNally EM, Sue E, Yantiss RK, Gross SS, Leve ND, Karoly ED, Suen C, Ling L. 2016. Celecoxib alters the intestinal microbiota and metabolome in association with reducing polyp burden. Cancer Prevention Research:canprevres. 0095.2016.

25. Smith EA, Macfarlane G. 1997. Dissimilatory amino acid metabolism in human colonic bacteria. Anaerobe 3:327–337.

26. Okuda S, Yamada T, Hamajima M, Itoh M, Katayama T, Bork P, Goto S, Kanehisa M. 2008. KEGG Atlas mapping for global analysis of metabolic pathways. Nucleic acids research 36:W423–W426.

27. Rautio M, Eerola E, Vaisanen-Tunkelrott M-L, Molitoris D. 2003. Reclassification of bacteroides putredinis (Weinberg et al., 1937) in a new genus alistipes gen. nov., as alistipes purtredinis comb. nov., and description of alistipes finegoldii sp. nov., from human sources. Systematic and applied microbiology 26:182.

28. Morotomi M, Nagai F, Sakon H, Tanaka R. 2008. Dialister succinatiphilus sp. nov. and Barnesiella intestinihominis sp. nov., isolated from human faeces. International journal of systematic and evolutionary microbiology 58:2716–2720.

29. Donohoe DR, Collins LB, Wali A, Bigler R, Sun W, Bultman SJ. 2012. The Warburg effect dictates the mechanism of butyrate-mediated histone acetylation and cell proliferation. Molecular cell 48:612–626.

30. Belcheva A, Irrazabal T, Robertson SJ, Streutker C, Maughan H, Rubino S, Moriyama EH, Copeland JK, Surendra A, Kumar S. 2014. Gut microbial metabolism drives transformation of MSH2-deficient colon epithelial cells. Cell 158:288–299.

31. Kang M, Martin A. Microbiome and colorectal cancer: Unraveling host-microbiota interactions in colitis-associated colorectal cancer development, p. In (ed), Elsevier,

32. Arber N, Eagle CJ, Spicak J, Rácz I, Dite P, Hajer J, Zavoral M, Lechuga MJ, Gerletti P, Tang J. 2006. Celecoxib for the prevention of colorectal adenomatous polyps. New England Journal of Medicine 355:885–895.

33. Umar A, Steele VE, Menter DG, Hawk ET. Mechanisms of nonsteroidal anti-inflammatory drugs in cancer prevention, p 65–77. In (ed), Elsevier,

34. Wasielewski H, Alcock J, Aktipis A. 2016. Resource conflict and cooperation between human host and gut microbiota: implications for nutrition and health. Annals of the New York Academy of Sciences 1372:20–28.

35. Röhrig F, Schulze A. 2016. The multifaceted roles of fatty acid synthesis in cancer. Nature Reviews Cancer 16:732.

36. Schug ZT, Voorde JV, Gottlieb E. 2016. The metabolic fate of acetate in cancer. Nature Reviews Cancer 16:708.

37. Schug ZT, Peck B, Jones DT, Zhang Q, Grosskurth S, Alam IS, Goodwin LM, Smethurst E, Mason S, Blyth K. 2015. Acetyl-CoA synthetase 2 promotes acetate utilization and maintains cancer cell growth under metabolic stress. Cancer cell 27:57–71.

38. Macfarlane S, Macfarlane GT. 2003. Regulation of short-chain fatty acid production. Proceedings of the Nutrition Society 62:67–72.

39. Wu W, Sun M, Chen F, Cao AT, Liu H, Zhao Y, Huang X, Xiao Y, Yao S, Zhao Q. 2017. Microbiota metabolite short-chain fatty acid acetate promotes intestinal IgA response to microbiota which is mediated by GPR43. Mucosal immunology 10:946.

40. Duncan SH, Barcenilla A, Stewart CS, Pryde SE, Flint HJ. 2002. Acetate utilization and butyryl coenzyme A (CoA): acetate-CoA transferase in butyrate-producing bacteria from the human large intestine. Applied and environmental microbiology 68:5186–5190.

41. Yutin N, Galperin MY. 2013. A genomic update on clostridial phylogeny: Gram-negative spore formers and other misplaced clostridia. Environmental microbiology 15:2631–2641.

42. Warnick TA, Methe BA, Leschine SB. 2002. Clostridium phytofermentans sp. nov., a cellulolytic mesophile from forest soil. International journal of systematic and evolutionary microbiology 52:1155–1160.

43. Gophna U, Konikoff T, Nielsen HB. 2017. Oscillospira and related bacteria–From metagenomic species to metabolic features. Environmental microbiology 19:835–841.

44. Mills S, Stanton C, Lane JA, Smith GJ, Ross RP. 2019. Precision nutrition and the microbiome, Part I: Current state of the science. Nutrients 11:923.

45. Louis P, Flint HJ. 2017. Formation of propionate and butyrate by the human colonic microbiota. Environmental microbiology 19:29–41.

46. Vital M, Howe AC, Tiedje JM. 2014. Revealing the bacterial butyrate synthesis pathways by analyzing (meta) genomic data. MBio 5:e00889–14.

47. Geirnaert A, Steyaert A, Eeckhaut V, Debruyne B, Arends JB, Van Immerseel F, Boon N, Van de Wiele T. 2014. Butyricicoccus pullicaecorum, a butyrate producer with probiotic potential, is intrinsically tolerant to stomach and small intestine conditions. Anaerobe 30:70–74.

48. Vital M, Gao J, Rizzo M, Harrison T, Tiedje JM. 2015. Diet is a major factor governing the fecal butyrate-producing community structure across Mammalia, Aves and Reptilia. The ISME journal 9:832.

49. Kee J-Y, Ito A, Hojo S, Hashimoto I, Igarashi Y, Tsuneyama K, Tsukada K, Irimura T, Shibahara N, Takasaki I. 2014. CXCL16 suppresses liver metastasis of colorectal cancer by promoting TNF-α-induced apoptosis by tumor-associated macrophages. BMC cancer 14:949.

50. De Weirdt R, Hernandez-Sanabria E, Fievez V, Mees E, Geirnaert A, Van Herreweghen F, Vilchez-Vargas R, Van den Abbeele P, Jauregui R, Pieper DH. 2017. Mucosa-associated biohydrogenating microbes protect the simulated colon microbiome from stress associated with high concentrations of poly-unsaturated fat. Environmental microbiology 19:722–739.

51. Truchado P, Hernandez-Sanabria E, Salden BN, Van den Abbeele P, Vilchez-Vargas R, Jauregui R, Pieper DH, Possemiers S, Van de Wiele T. 2017. Long chain arabinoxylans shift the mucosa-associated microbiota in the proximal colon of the simulator of the human intestinal microbial ecosystem (M-SHIME). Journal of Functional Foods 32:226–237.

52. El Hage R, Hernandez-Sanabria E, Calatayud Arroyo M, Props R, Van de Wiele T. 2019. Propionate-producing consortium restores antibiotic-induced dysbiosis in a dynamic in vitro model of the human intestinal microbial ecosystem. Frontiers in Microbiology 10:1206.

53. Hernandez-Sanabria E, Goonewardene LA, Wang Z, Zhou M, Moore SS. 2013. Influence of sire breed on the interplay among rumen microbial populations inhabiting the rumen liquid of the progeny in beef cattle. PloS one 8:e58461.

54. McMurdie PJ, Holmes S. 2013. phyloseq: an R package for reproducible interactive analysis and graphics of microbiome census data. PloS one 8:e61217.

55. McMurdie PJ, Holmes S. 2014. Waste not, want not: why rarefying microbiome data is inadmissible. PLoS computational biology 10:e1003531.

56. Grunert O, Hernandez-Sanabria E, Vilchez-Vargas R, Jauregui R, Pieper DH, Perneel M, Van Labeke M-C, Reheul D, Boon N. 2016. Mineral and organic growing media have distinct community structure, stability and functionality in soilless culture systems. Scientific reports 6:18837.

57. Oksanen J, Blanchet FG, Kindt R, Legendre P, Minchin PR, O’hara R, Simpson GL, Solymos P, Stevens MHH, Wagner H. 2013. Package ‘vegan’. Community ecology package, version 2.

58. Kurtz ZD, Müller CL, Miraldi ER, Littman DR, Blaser MJ, Bonneau RA. 2015. Sparse and compositionally robust inference of microbial ecological networks. PLoS computational biology 11:e1004226.

59. Bastian M, Heymann S, Jacomy M. 2009. Gephi: an open source software for exploring and manipulating networks. Icwsm 8:361–362.

60. Lê Cao K-A, Boitard S, Besse P. 2011. Sparse PLS discriminant analysis: biologically relevant feature selection and graphical displays for multiclass problems. BMC bioinformatics 12:253.

61. Paul W, Marta C. 2018. Resolving host–microbe interactions in the gut: the promise of in vitro models to complement in vivo research. Current opinion in microbiology 44:28–33.

62. Calatayud MA, de Wiele Van T, Hernandez-Sanabria E. 2018. Assessing the Viability of a Synthetic Bacterial Consortium on the In Vitro Gut Host-microbe Interface. Journal of visualized experiments: JoVE.

