## Supplementary Information for "Short term supplementation of celecoxib shifted butyrate production on a simulated model of the gut microbial ecosystem and ameliorated *in vitro* inflammation"

Running title: Celecoxib shifts butyrate in an *in vitro* gut microbial ecosystem

**Illumina library preparation and purification**

The V5-V6 hypervariable region of the 16S rRNA gene from the cDNA was amplified using primers 341F and 785R. Illumina sequencing adapters and dual-index barcodes were added to the amplicon, using a limited-cycle PCR that included an initial denaturation step at 95 ºC for 3 min, 15 cycles of a denaturation step at 95ºC for 30 s, an annealing step at 55 ºC for 10s, an extension step at 72 ºC for 45 s, and a final extension at 72 ºC for 5 min. Following, a clean-up step was performed using the AMPure XP beads (Beckman-Coulter, Krefeld, Germany) to remove free primers and primer-dimer species from amplicons. A second PCR to attach the specific Illumina multiplexing sequencing primers and index primers, was performed. Thermal cycling included an initial denaturation step at 95 ºC for 3 min, 8 cycles of a denaturation step at 95ºC for 30 s, an annealing step at 55 ºC for 30 s, an extension step at 72 ºC for 30 s, and a final extension at 72 ºC for 5 min.

PCR products were verified by gel electrophoresis, purified using the Promega Wizard PCR clean-up kit (Promega, Madison, WI, USA) following the manufacturer's instructions and quantified with the QuantiFluor dsDNA System kit (Promega, Leiden, The Netherlands). High-throughput amplicon sequencing was performed with the Illumina MiSeq platform according to the manufacturer's guidelines at LGC Genomics GmbH (Berlin, Germany).

Contigs were created by merging paired-end reads based on the Phred quality score (of both reads) heuristic as described by (17) in Mothur v.1.33.3 (18). Contigs were aligned to the SILVA database and filtered from those with (i) ambiguous bases, (ii) more than 10 homopolymers, and (iii) those not corresponding to the targeted region. Chimera removal and operational taxonomic unit (OTU) clustering of the sequencing reads was performed using UCHIME, with the nearest neighbour clustering algorithm implemented in mothur, at 0.03 distance (19). Phylotype representatives were then generated by clustering at 97% similarity (1 mismatch), with a confidence level of at least 80 with Cyanobacteria, Eukaryota, and Archaea lineages removed. For taxonomic classification, sequence composition of the dataset was compared using the RDP Classifier tool (20), and the RDP trainset version 9. Quality of the sequencing and post-processing pipeline was verified by incorporating mock samples (n = 12 species) in triplicate into the same sequencing run. After examining read counts, if any OTU was not classified up to genus level, the consensus sequence was blasted using the SILVA database (21) to obtain an approximate taxonomic classification. Singletons that remained unclassified were culled.

Supplementary Table 1A. Mean starting relative percentages of the three major SCFA derived from microbial fermentation, following supplementation with either celecoxib (CX) or the carrier (PEG). N = 6, means between treatments were not significantly different.

| SCFA (%) | Ratio found in colon and stool Cummings J.H et al. (1987) | Donor 3 | | Donor 4 | | Donor 5 | | Donor 6 | | Donor 7 | | Donor 8 | |
|---|---|---|---|---|---|---|---|---|---|---|---|---|---|
| | | CX | PEG | CX | PEG | CX | PEG | CX | PEG | CX | PEG | CX | PEG |
| Acetate | 60 | 60.02 | 65.71 | 35.45 | 39.30 | 63.90 | 65.32 | 59.25 | 63.42 | 40.88 | 41.84 | 49.98 | 53.97 |
| Propionate | 20 | 23.20 | 21.64 | 14.84 | 14.46 | 12.63 | 12.48 | 12.83 | 11.70 | 14.20 | 13.97 | 10.61 | 10.81 |
| Butyrate | 20 | 15.49 | 12.65 | 10.17 | 9.48 | 15.85 | 15.52 | 8.20 | 5.84 | 6.92 | 7.20 | 6.23 | 6.15 |

Supplementary Table 1B. Relative percentages of the three major SCFA derived from microbial fermentation, following supplementation with either celecoxib (CX) or the carrier (PEG) and incubated for 16 h. N = 8; **, *P* < 0.05. ***, *P* < 0.0001

| SCFA (%) | Ratio found in colon and stool Cummings J.H et al. (1987) | Donor 1 | | Donor 2 | | Donor 3 | | Donor 4 | | Donor 5 | | Donor 6 | | Donor 7 | | Donor 8 | |
|---|---|---|---|---|---|---|---|---|---|---|---|---|---|---|---|---|---|
| | | CX | PEG | CX | PEG | CX | PEG | CX | PEG | CX | PEG | CX | PEG | CX | PEG | CX | PEG |
| Acetate | 60 | 49.41 | 47.15 | 74.64 | 62.51** | 22.46 | 28.97 | 33.32 | 34.76 | 73.82 | 57.87*** | 84.89 | 63.64*** | 68.12 | 67.87 | 73.99 | 69.94 |
| Propionate | 20 | 17.30 | 18.05 | 22.40 | 18.27 | 21.01 | 18.89 | 6.90 | 9.06 | 14.96 | 16.13 | 12.58 | 18.93** | 16.14 | 13.88 | 14.18 | 13.48 |
| Butyrate | 20 | 15.13 | 17.15 | 2.27 | 17.89*** | 4.83 | 11.76*** | 7.51 | 7.89 | 6.04 | 19.48*** | 2.46 | 13.01*** | 8.15 | 10.29** | 7.66 | 8.07 |

Supplementary Table 2. Total metabolically active bacterial population at the beginning (A) and at after 16h (B) of incubation with either celecoxib (CX) or carrier (PEG). Means presented were calculated from 3 replicate incubations run in triplicate per donor. N = 8, NS = not significantly different.

| *0 h* | | | | | *16 h* | | | |
|---|---|---|---|---|---|---|---|---|
| Donor | Treatment | | SEM | *P* value | Treatment | | SEM | *P* value |
| | CX | PEG | | | CX | PEG | | |
| D1 | - | - | - | - | 9.78E+09 | 2.11E+10 | | <0.001 |
| D2 | - | - | - | - | 4.90E+09 | 4.72E+09 | | NS |
| D3 | 1.54E+08 | 4.34E+07 | 4.11E+07 | NS | 8.59E+08 | 2.86E+08 | | NS |
| D4 | 1.49E+09 | 1.20E+09 | 3.88E+08 | NS | 1.79E+09 | 1.71E+09 | 9.41E+08 | NS |
| D5 | 3.11E+09 | 9.08E+09 | 2.54E+09 | NS | 3.00E+09 | 4.73E+09 | | NS |
| D6 | 5.70E+08 | 3.47E+09 | 4.25E+08 | 0.0007 | 2.49E+08 | 2.78E+08 | | NS |
| D7 | 2.32E+08 | 3.43E+08 | 7.87E+07 | NS | 1.08E+09 | 1.64E+09 | | NS |
| D8 | 6.53E+08 | 1.12E+09 | 1.36E+08 | 0.04 | 5.30E+09 | 3.21E+08 | | 0.04 |

Supplementary Table 3. Community diversity metrics using the Hill diversity of order 2, or inverse Simpson index ($D_2$) after exposure to celecoxib (CX) or the carrier (PEG), n = 3

| Donor | Time | Inverse Simpson | | SEM | *P* value | | |
|---|---|---|---|---|---|---|---|
| | | CX | PEG | | Time | Treatment | Time*Treatment |
| D1 | 0h | - | - | 4.41 | - | 0.59 | - |
| | 16h | 7.20 | 11.16 | | | | |
| D2 | 0h | - | - | 1.85 | - | 0.56 | - |
| | 16h | 6.08 | 7.89 | | | | |
| D3 | 0h | 3.32 | 4.33 | 0.42 | 0.006 | 0.01 | 0.04 |
| | 16h | 3.90 | 7.43 | | | | |
| D4 | 0h | 6.93 | 9.13 | 0.60 | 0.04 | 0.03 | 0.003 |
| | 16h | 12.80 | 7.14 | | | | |
| D5 | 0h | 10.42 | 9.51 | 1.16 | 0.70 | 0.04 | 0.73 |
| | 16h | 6.54 | 6.48 | | | | |
| D6 | 0h | 5.41 | 4.16 | 0.58 | 0.66 | 0.01 | 0.17 |
| | 16h | 6.81 | 7.50 | | | | |
| D7 | 0h | 8.86 | 9.99 | 0.99 | 0.23 | 0.01 | 0.79 |
| | 16h | 4.57 | 6.26 | | | | |
| D8 | 0h | 13.11 | 13.76 | 1.79 | 0.80 | 0.47 | 0.55 |
| | 16h | 12.85 | 11.20 | | | | |

Supplementary Table 4. Impact of donor, treatment and time on bacterial relative abundances following short-term supplementation of either celecoxib or the carrier. CX, celecoxib. PEG, carrier. Tpt, time point effect; Trt, treatment effect. N = 8.

| Taxa | Time point | CX | | PEG | | *P* value | | | | |
|---|---|---|---|---|---|---|---|---|---|---|
| | | Mean | SEM | Mean | SEM | Donor | Tpt | Trt | Trt*Tpt interaction | Donor*Trt*Tpt interaction |
| *Allisonella* | 0h | 0.0001 | 0.00003 | 0.0002 | 0.0001 | 0.02 | 0.07 | 0.40 | 0.52 | 0.52 |
| | 16h | 0.0002 | 0.0001 | 0.0003 | 0.0001 | | | | | |
| *Anaerostipes* | 0h | 0.0015 | 0.0004 | 0.0015 | 0.0003 | 0.002 | 0.11 | 0.31 | 0.40 | 0.001 |
| | 16h | 0.0008 | 0.0002 | 0.0013 | 0.0002 | | | | | |
| *Coprococcus* 2 | 0h | 0.0016 | 0.0005 | 0.0010 | 0.0003 | <0.0001 | 0.20 | 0.63 | 0.30 | 0.02 |
| | 16h | 0.0008 | 0.0003 | 0.0010 | 0.0002 | | | | | |
| Erysipelotrichaceae UCG 003 | 0h | 0.0017 | 0.0003 | 0.0014 | 0.0002 | 0.003 | 0.16 | 0.11 | 0.46 | <0.0001 |
| | 16h | 0.0015 | 0.0003 | 0.001 | 0.0002 | | | | | |
| *Flavonifractor* | 0h | 0.0003 | 0.0001 | 0.0002 | 0.0001 | 0.001 | 0.15 | 0.43 | 0.08 | 0.0004 |
| | 16h | 0.0001 | 0.00003 | 0.0002 | 0.0001 | | | | | |
| *Fusobacterium* | 0h | 0.0016 | 0.0005 | 0.0012 | 0.0004 | <0.0001 | 0.85 | 0.65 | 0.63 | 0.66 |
| | 16h | 0.0015 | 0.0005 | 0.0015 | 0.0004 | | | | | |
| Lachnospiraceae FCS020 group | 0h | 0.0001 | 0.00004 | 0.0002 | 0.0001 | 0.01 | 0.16 | 0.13 | 0.79 | 0.70 |
| | 16h | 0.0001 | 0.00002 | 0.0001 | 0.00003 | | | | | |
| Lachnospiraceae UCG 001 | 0h | 0.0012 | 0.0003 | 0.0011 | 0.0002 | 0.11 | 0.004 | 0.99 | 0.63 | 0.24 |
| | 16h | 0.0004 | 0.0002 | 0.0005 | 0.0001 | | | | | |
| *Marvinbryantia* | 0h | 0.0001 | 0.00003 | 0.00004 | 0.00001 | 0.17 | <0.0001 | 0.87 | 0.11 | 0.43 |
| | 16h | 0.000004 | 0.000001 | 0.00001 | 0.000002 | | | | | |

| Taxa | Time point | CX | | PEG | | *P* value | | | | |
|---|---|---|---|---|---|---|---|---|---|---|
| | | Mean | SEM | Mean | SEM | Donor | Tpt | Trt | Trt*Tpt interaction | Donor*Trt*Tpt interaction |
| *Parabacteroides* | 0h<br>16h | 0.0095<br>0.0109 | 0.0017<br>0.0014 | 0.0075<br>0.0080 | 0.0014<br>0.0009 | <0.0001 | 0.54 | 0.11 | 0.82 | <0.0001 |
| *Paraprevotella* | 0h<br>16h | 0.0005<br>0.0003 | 0.0002<br>0.0001 | 0.0005<br>0.0004 | 0.0002<br>0.0001 | 0.0001 | 0.13 | 0.65 | 0.77 | 0.02 |
| *Parasutterella* | 0h<br>16h | 0.0013<br>0.0020 | 0.0003<br>0.0004 | 0.0013<br>0.0013 | 0.0003<br>0.0003 | <0.0001 | 0.33 | 0.46 | 0.36 | 0.02 |
| *Peptococcus* | 0h<br>16h | 0.0003<br>0.0001 | 0.0001<br>0.00004 | 0.0001<br>0.0001 | 0.00004<br>0.00003 | 0.01 | 0.06 | 0.25 | 0.41 | 0.68 |
| *Phascolarctobacterium* | 0h<br>16h | 0.0035<br>0.0027 | 0.0008<br>0.0006 | 0.0028<br>0.0029 | 0.0006<br>0.0005 | <0.0001 | 0.56 | 0.77 | 0.53 | <0.0001 |
| *Ruminiclostridium* 5 | 0h<br>16h | 0.0006<br>0.0003 | 0.0001<br>0.0001 | 0.0005<br>0.0003 | 0.0001<br>0.0001 | 0.09 | 0.02 | 0.49 | 0.83 | 0.03 |
| *Ruminiclostridium* 6 | 0h<br>16h | 0.0013<br>0.0007 | 0.0003<br>0.0002 | 0.0012<br>0.0009 | 0.0003<br>0.0002 | <0.0001 | 0.09 | 0.80 | 0.54 | 0.02 |
| Ruminococcaceae NK4A214 group | 0h<br>16h | 0.0004<br>0.0002 | 0.0001<br>0.0001 | 0.0003<br>0.0002 | 0.0001<br>0.0001 | <0.0001 | 0.10 | 0.57 | 0.49 | 0.28 |
| Ruminococcaceae UCG 004 | 0h<br>16h | 0.0002<br>0.0001 | 0.0001<br>0.00003 | 0.0002<br>0.0001 | 0.0001<br>0.00003 | 0.001 | 0.06 | 0.73 | 0.60 | 0.75 |
| Ruminococcaceae UCG 010 | 0h<br>16h | 0.0004<br>0.0002 | 0.0001<br>0.0001 | 0.0003<br>0.0002 | 0.0001<br>0.0001 | 0.0004 | 0.07 | 0.95 | 0.46 | 0.03 |
| *Ruminococcus* 2 | 0h<br>16h | 0.0017<br>0.0014 | 0.0005<br>0.0004 | 0.0015<br>0.0011 | 0.0003<br>0.0002 | <0.0001 | 0.33 | 0.49 | 0.83 | 0.01 |

| Taxa | Time point | CX | | PEG | | *P* value | | | | |
|---|---|---|---|---|---|---|---|---|---|---|
| | | Mean | SEM | Mean | SEM | Donor | Tpt | Trt | Trt*Tpt interaction | Donor*Trt*Tpt interaction |
| *Streptococcus* | 0h<br>16h | 0.00002<br>0.00005 | 0.00001<br>0.00002 | 0.00002<br>0.00003 | 0.00001<br>0.00001 | 0.01 | 0.06 | 0.79 | 0.23 | 0.17 |
| Unclassified Christensenellaceae | 0h<br>16h | 0.0001<br>0.00004 | 0.00002<br>0.00001 | 0.00003<br>0.00004 | 0.00001<br>0.00001 | 0.002 | 0.79 | 0.32 | 0.21 | 0.79 |
| *Veillonella* | 0h<br>16h | 0.0009<br>0.0013 | 0.0003<br>0.0003 | 0.0007<br>0.0009 | 0.0002<br>0.0003 | 0.03 | 0.42 | 0.34 | 0.80 | 0.01 |
| *Victivallis* | 0h<br>16h | 0.0008<br>0.0005 | 0.0002<br>0.0001 | 0.0004<br>0.0004 | 0.0001<br>0.0001 | <0.0001 | 0.46 | 0.09 | 0.30 | 0.01 |
